# MicroRNA-511-3p mediated modulation of the peroxisome proliferator-activated receptor gamma (PPARγ) controls LPS-induced inflammatory responses in human monocyte derived DCs

**DOI:** 10.1101/2020.11.05.369967

**Authors:** Dennis Awuah, Alisa Ruisinger, Meshal Alobaid, Chidimma Mbadugha, Amir M. Ghaemmaghami

**Affiliations:** Division of Immunology, School of Life Sciences, Faculty of Medicine and Health Sciences, University of Nottingham, Nottingham, NG7 2RD, United Kingdom; T cell Therapeutics Research Laboratory, Beckman Research Institute, City of Hope, Duarte, CA, 91010, United States

**Author notes:** Correspondence address and reprint request to Professor Amir M. Ghaemmaghami, Immunology and Immuno-bioengineering group, School of Life Sciences, Faculty of Medicine and Health Sciences, University Park, University of Nottingham, Nottingham NG7 2RD, UK.

**Keywords:** Dendritic cells, miR-511-3p, RNAi, PPARγ, Inflammation, Indoleamine 2,3 dioxygenase, Immune modulation

## Abstract

The peroxisome proliferator activated receptor gamma (PPARγ) is a ligand activated transcription factor expressed in dendritic cells (DCs), where it exerts anti-inflammatory responses against TLR4-induced inflammation. Recently, microRNA-511 (miR-511) has also emerged as a key player in controlling TLR4-mediated signalling, and in regulating the function of DCs. Interestingly, PPARγ has been previously highlighted as a putative target of miR-511 activity; however the link between miR-511 and PPARγ and its influence on human DC function within the context of LPS-induced inflammatory responses is unknown. Using a selection of miR-511-3p-specific inhibitors and mimics, we demonstrate for the first time that up or downregulation of miR-511-3p inversely correlates with PPARγ mRNA levels and transcriptional activity following treatment with PPARγ synthetic agonist rosiglitazone (RSG), in the presence or absence of LPS. Additionally, we show that PPARγ activation with RSG modulates LPS-induced DC activation and downregulates pro-inflammatory cytokine production following downregulation of miR-511-3p. Lastly, PPARγ activation was shown to suppress LPS-mediated induction of indoleamine 2,3-dioxygenase (IDO) activity in DCs, most likely due to changes in miR-511-3p expression. These data suggest that PPARγ-induced modulation of DC phenotype and function is influenced by miR-511-3p expression, which may serve as a potential therapeutic target against inflammatory diseases.

## Introduction

The innate immune system rapidly responds to endotoxin exposure through the induction of acute inflammatory responses (1). Endotoxins such as LPS, a major cell wall component of Gram-negative bacteria, can induce potent inflammatory responses through the release of an array of mediators such as cytokines, chemokines and growth factors. However, when uncontrolled, inflammatory responses can lead to tissue damage and chronic inflammatory diseases (2). Dendritic cells (DCs) are the most efficient antigen presenting cells (APCs) capable of initiating and maintaining primary immune responses and are key players in regulating inflammatory responses (3). DCs are highly sensitive to even low concentrations of LPS in the environment and detect the presence of LPS (for instance during bacterial infection) via Toll-like receptor 4 (TLR4). This triggers downstream signaling pathways, which co-ordinate expression of genes required to initiate or control inflammation (4).

Recently, the role of the peroxisome proliferator-activated receptor γ (PPARγ) in modulating inflammatory responses has been of particular interest. PPARγ is a lipid-activated transcription factor expressed in a variety of cell types including DCs, where it regulates genes associated with adipogenesis, lipid metabolism and inflammation (5). In DCs, PPARγ regulates other various processes including maturation and migration, activation, antigen presentation and cytokine production (6–8). Moreover, PPARγ activation in DCs in the presence of LPS resulted in decreased expression of the pro-inflammatory cytokine IL-12 (9), suggesting that PPARγ ligands may promote anti-inflammatory responses. Studies by us and others have also demonstrated a key role for the tryptophan metabolizing enzyme, IDO in controlling LPS-mediated inflammatory responses (10–13). Interestingly, the recent finding that PPARγ promotes IDO activity and subsequent immune suppression in the tumour microenvironment highlights the complexity of PPARγ-induced regulatory mechanisms (14). Nevertheless, the overall immunosuppressive effect of PPARγ and the underlying mechanisms particularly during LPS-induced inflammation in DCs are not fully understood.

MicroRNAs (miRNAs) are a class of gene regulators that bind the 3’UTR of target genes and cause translational inhibition or mRNA degradation (15). Due to their role in regulating different biological processes, miRNAs represent crucial regulators in human health and disease. Recently, miR-511-3p (located at the ‘3 end of the mature miR-511 strand) has been identified as an important regulator of human DC and macrophage development and function. For example, miR-511 was shown to be a key regulator of TLR4 expression in human DCs (16). Additionally, transcriptional products associated with inflammation and wound healing in alternatively activated macrophage (AAM), were found to be altered by miR-511 expression (17). Interestingly, PPARγ has also been highlighted as a putative target of miR-511 activity, affecting human myeloid cell differentiation and function (16,18) however the mechanisms underlying these events are not yet known. Here, we sought to investigate the potential link between miR-511-3p expression and PPARγ activity and how this influences LPS-mediated inflammatory responses in human DCs. Better understanding of the mechanisms regulating PPARγ activity could pave the way for rational design of therapies against a range of inflammatory disorders.

## Materials and Methods

### Generation of monocyte-derived dendritic cells (DCs)

This was done as previously described (19,20). Buffy coats were obtained from healthy donors (National Blood Service, Sheffield, UK) after obtaining informed written consent and following ethics committee approval (Research Ethics Committee, Faculty of Medicine and Health Sciences, University of Nottingham). All methods were performed in accordance with the relevant guidelines and regulations. PBMCs were separated by density gradient centrifugation on Histopaque (Sigma-Aldrich, UK). Monocytes were purified by positive selection using the MACS CD14 isolation kit (Miltenyi Biotec) and cultured in 24 well plates, using RPMI medium supplemented with 10% heat-inactivated FBS, 100U/ml penicillin, 100U/ml streptomycin and 2mM L-glutamine (all from Sigma). Cells were incubated at 37 °C with 5% CO_2_ in a humidified incubator. The purity of CD14^+^ cells was always above 90% as measured by flow cytometry. DC differentiation was carried out over 6 days with 250 U/ml IL-4 and 50ng/ml GM-CSF (Miltenyi Biotec).

### Flow cytometry analysis

Mouse monoclonal antibodies against human CD86 (clone FM95) and DC-SIGN (clone DCN47.5) were purchased from Miltenyi Biotec. Antibodies against human CD83 (clone HB15e) was purchased from eBioscience. Anti-PDL1 (clone MIH1) antibody was purchased from BD Biosciences (San Jose, Calif.) and the anti-CD206 antibody (clone 15-2) was purchased from Biolegend. Briefly, cells were collected and washed twice in cold PBA (PBS buffer containing 0.5% BSA and 0.1% sodium azide (Sigma Aldrich)). Staining with labelled antibodies was then carried out in the dark at 4 °C for 20 minutes according to manufacturer’s instructions. Unless otherwise stated, specific antibodies were conjugated to Fluorescein Isothiocyanate (FITC), Phycoerythrin (PE) or Phycoerythrin Cyanine 5.1 (PE/Cy5). Following staining, samples were washed twice with PBA buffer and fixed in 0.5% formaldehyde solution before analysis. Non-reactive, isotype-matched antibodies were used as controls. Flow cytometry was carried out on the FC500 Flow Cytometer (Beckman Coulter, UK) and data was analysed using Weasel Software for Windows.

### Quantification of IDO activity

IDO activity was determined as described before (19,21). Briefly, DCs (2.5~10^5^cells/ml) were seeded in a 24 well plate with complete RPMI media, supplemented with 100μM L-tryptophan (TRP) (Sigma Aldrich). Cells were then stimulated with the PPARγ agonist and antagonist RSG and GW9662 (5μM) respectively, in the presence and absence of LPS (0.1μg/ml). A colorimetric assay for IDO activity was performed after 24 hours of stimulation by measuring the levels of L-kynurenine (KYN) produced in culture supernatant. The concentration of L-KYN was then calculated from a standard curve of defined concentrations from 0 to 200μM. Rosiglitazone (RSG) and GW9662 were purchased from Cayman chemicals (Cambridge, UK) and L-KYN was obtained from Sigma Aldrich, UK.

### Live/Dead assay

Cell viability following treatment with RSG and GW9662 was determined using the LIVE/DEAD assay kit (Thermo Fisher, UK) as described previously (22). Cells were imaged using the Etaluma LS720 Microscope (Carlsbad, CA). Data was analysed using image J software.

### RNA interference

Pre-designed miR-511-3p inhibitors and mimics were purchased from Qiagen and transfection was carried out using the HiPerFect Transfection Reagent according to manufacturer’s protocol (Qiagen). The miRNA targeted sequence was 5’-AAUGUGUAGCAAAAGACAGA-3’. Briefly, CD14^+^ monocytes were suspended in Opti-MEM^®^ reduced serum media (Gibco) and seeded. Prior to transfection, miR-511-3p inhibitor or mimic was diluted in serum-free media with transfection reagent in separate tubes for 10 minutes at room temperature, before adding drop-wise unto cells. The miScript Inhibitor Negative Control and the AllStars Negative Control siRNA (Qiagen) were used as scrambled controls (CT) for inhibitor and mimic respectively. All transfections were carried out at a final concentration of 50nM as determined during optimizations. Monocytes were differentiated into DCs after 6 hours of incubation, with fresh Opti-MEM^®^ media supplemented with IL-4 and GM-CSF. Transfection efficiency was determined using the fluorescently-labelled siGLO RISC-free control siRNA (GE Healthcare) and miRNA and/or mRNA expression was assessed afterwards.

### RNA isolation and cDNA synthesis

Dendritic cell samples were washed twice with cold PBS and stored in RNA later before RNA isolation. miR-511-3p transfected cells as well as controls were stimulated on day 6 with RSG in the presence or absence of LPS (Sigma Aldrich) for 24 hours before harvesting. Total RNA, including small RNAs was isolated with Trizol Reagent using the miRNeasy Mini kit (Qiagen). The concentration of purified RNA was measured using a NanoDrop 1000 spectrophotometer (Thermo Scientific, Pierce). First-strand cDNA was then generated with the SuperScript III Reverse Transcription kit (Thermo Fisher Scientific) or the miScript Reverse Transcription (RT) II kit (Qiagen) or according to manufacturer’s instructions.

Reverse transcription was done with the T100 Thermal Cycler (Bio-Rad, UK) and the 5X HiFlex buffer was used for parallel qRT-PCR quantification of mature miRNA and mRNA.

### Real Time PCR (qRT-PCR)

Comparative real-time PCR for miR-511-3p expression was carried out on the MxPro 3005P qRT-PCR system, (Stratagene, CA) using the miScript SYBR Green PCR kit (Qiagen) according to manufacturer’s instructions. Briefly, 12.5μl 2X QuantiTect SYBR-Green Master mix, 2.5μl 10X Universal Primer and 2.5μl Primer Assay (forward primer) was mixed with 3ng of reverse transcription product. qRT-PCR cycling was initiated at 95 °C for 15 minutes, followed by 40 cycles of 94°C for 15 seconds, 55 °C for 30 seconds and 70 °C for 30 seconds. Mature miR-511-3p-specific primers were obtained from Qiagen and relative expression was normalised to U6 (RNU6-2) small nuclear RNAs. qRT-PCR for mRNA expression was done with the Brilliant III Ultra-Fast SYBR Green qRT-PCR Master Mix (Agilent Technologies) as previously described (19). The two-step cycling reaction was initiated at 95 °C for 3 minutes followed by 40 cycles of 95 °C for 20 seconds and 60 °C for 20 seconds. PPARG and FABP4 mRNA expression levels were normalised to GAPDH and calculated using the comparative Delta *C*t method. All experimental procedures were done in triplicate. Forward and reverse mRNA primers were selected using Roche Universal Probes Library and purchased from Eurofins Scientific, UK (Table 1)

**Table 1:**
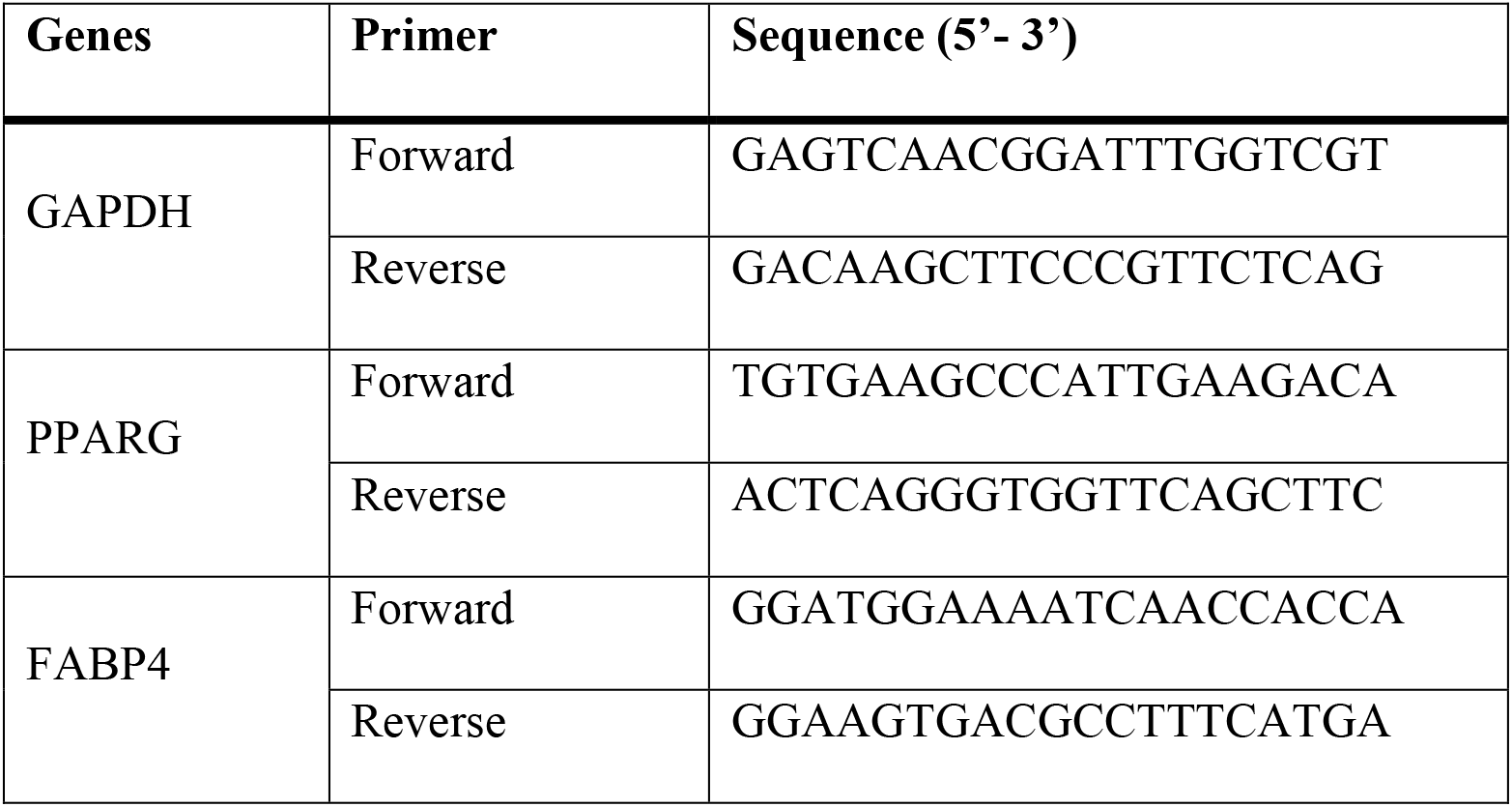
Primers for real-time PCR

### Cytokine ELISA

Culture supernatants were collected and stored at −20°C before analysis. The levels of TNF-α, IL-10 and IL-6 were measured by sandwich ELISA using the Duo Set ELISA kit (R&D Systems, UK) (23) according to manufacturer’s instructions.

### Statistical Analysis

Data were analysed using GraphPad Prism v7.02 for Windows (GraphPad Software, CA) and values expressed as mean ± standard deviation (SD) from three independent experiments unless otherwise stated. Statistical differences were determined using the student *t* test (to compare two groups) or one/two-way ANOVA (to compare three or more groups) with Tukey’s post-hoc testing. A *p* value <0.05 was considered statistically significant.

## Results

### Changes in miR-511-3p expression affect PPARγ expression and activity in human DCs

We first examined PPARγ regulation in human DC after changes in miR-511-3p expression. For this purpose, we differentiated monocyte-derived DCs in the presence of miR-511-3p-specific inhibitors and mimics, which resulted in a significant downregulation and/or increase in miRNA expression respectively (Figure 1). Subsequently, we measured the expression of PPARγ by comparative qRT-PCR and show for the first time that knockdown of miR-511-3p (henceforth referred to as miR-511-3p^low^), resulted in a significant increase in PPARγ mRNA levels (Figure 1A); whereas a significant decrease in PPARγ expression was seen in miR-511-3p overexpressed cells (i.e. miR-511-3p^hi^) (Figure 1B). As demonstrated in Figure 1C, there is a strong, inverse association between the two conditions, (i.e. the level of PPARγ relative to miR-511-3p) indicated by an increase in PPARγ expression when miR-511-3p is low and vice versa, when miR-511-3p expression is high. It is important to note that transfection with either miR-511-3p inhibitors or mimics did not result in loss of cell viability as determined by Annexin-V staining (Figure S1) and did not impact monocyte to DC differentiation as we have previously demonstrated (24).

**Figure 1:**
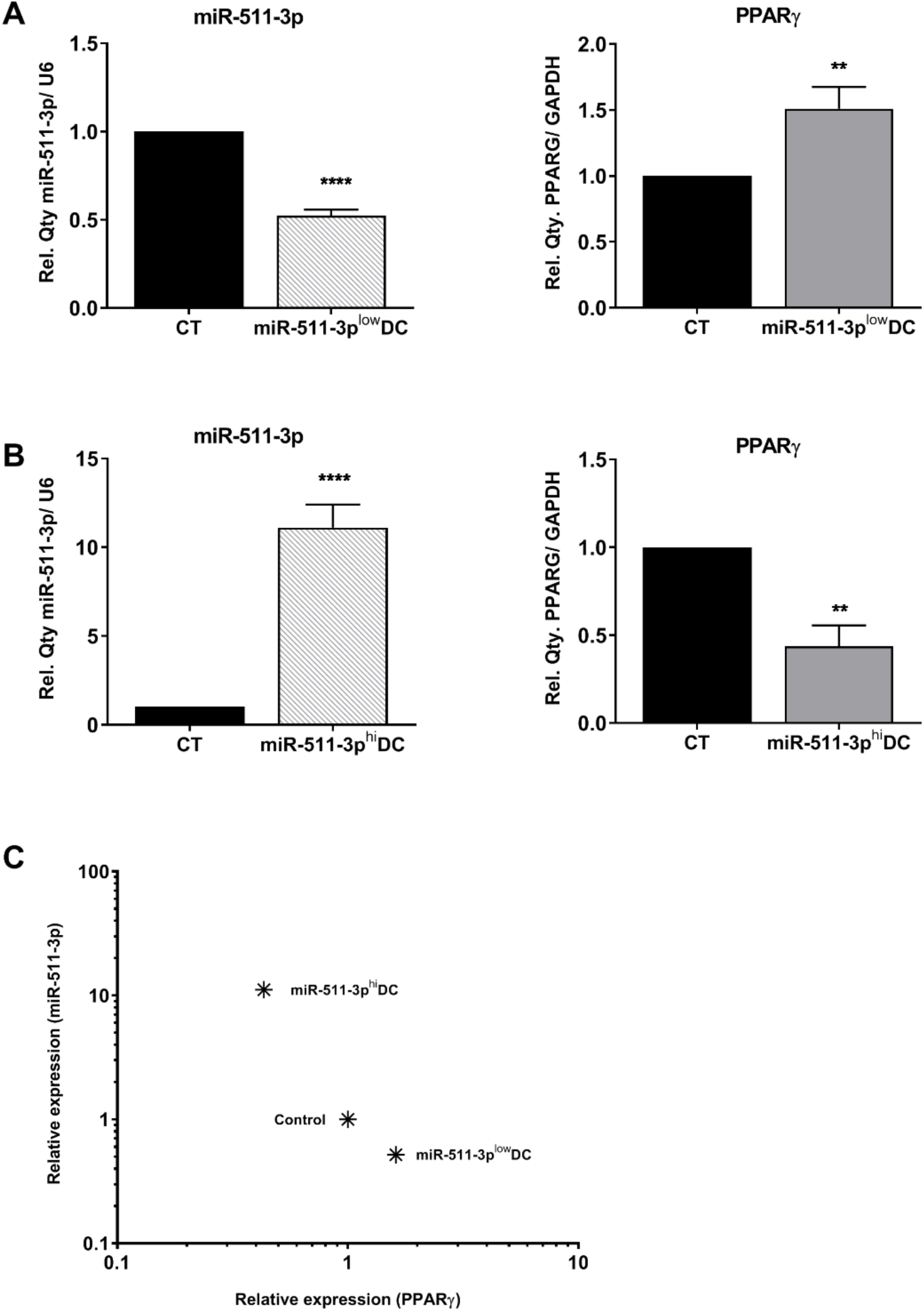
Analysis of PPARγ mRNA levels in response to up or downregulation of miR-511-3p expression. Relative expression of miR-511-3p and PPARγ respectively in A) miR-511-3p^low^ and B) miR-511-3p^hi^ DCs. C) Logarithmic scatter plot showing relationship between miR-511-3p and PPARγ expression. Monocytes were differentiated in the presence of 50nM miR-511-3p-specific inhibitors and mimics. Gene expression was assessed on day 6 and normalised to U6 and GAPDH for miRNA and mRNA respectively. (One representative data is shown out of three). **p < 0.01, ****p < 0.0001.

A number of natural (e.g. 15d-PGJ_2_) as well as synthetic agonists (also known as thiazolidinediones - TZDs) have been shown to induce PPARγ activation in a variety of cell types (25,26). Following up or downregulation of miR-511-3p and its effect on PPARγ expression, we treated DCs with LPS and the synthetic PPARγ agonist rosiglitazone (RSG) in order to determine the influence of these ligands on PPARγ activity in these cells. PPARγ activity was assessed by measuring the expression of one of its target genes (i.e. *FABP4*) by qRT-PCR. Interestingly, we found that PPARγ activity was significantly increased in miR-511-3p^low^ DCs compared to scrambled controls (CT), following RSG stimulation alone and in the presence of LPS (RSG+LPS) (Figure 2A). In contrast, we found a significant decrease in PPARγ activity in miR-511-3p^hi^ DCs treated with RSG+LPS but not with RGS only (Figure 2B).

**Figure 2:**
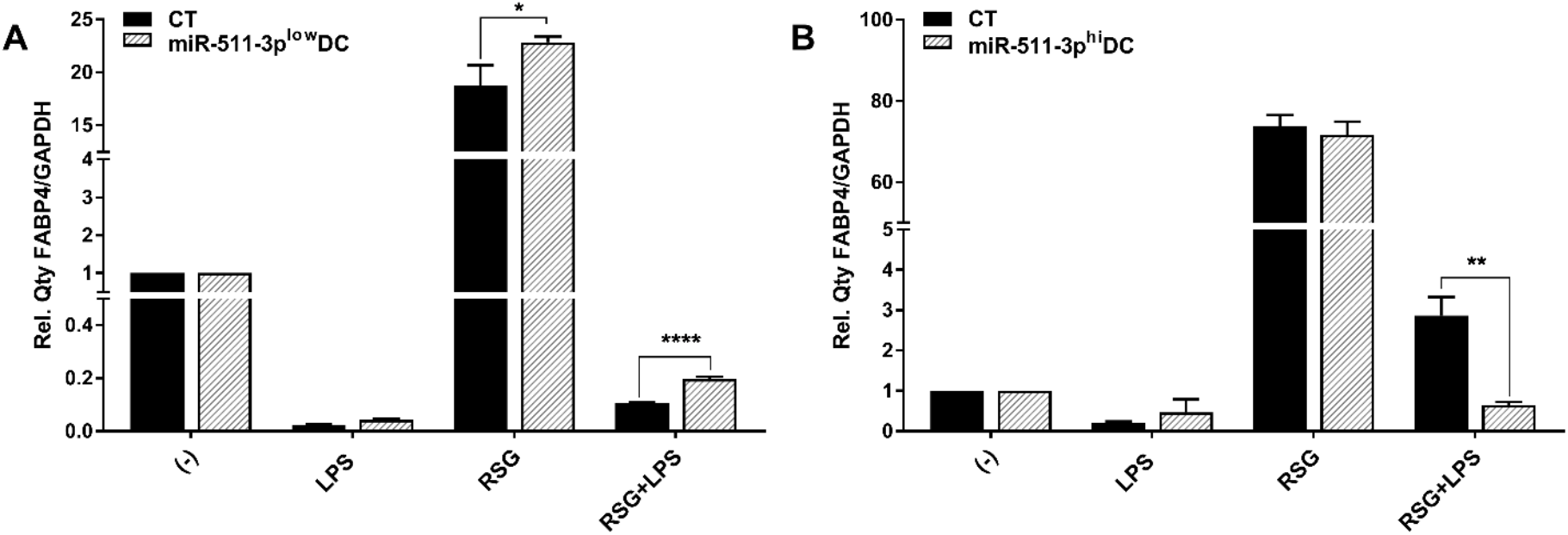
PPARγ activity is affected by changes in miR-511-3p expression. Relative *FABP4* expression in A) miR-511-3p^low^ and B) miR-511-3p^hi^ DCs treated with RSG and LPS. Monocyte derived DCs were transfected with 50nM inhibitor and mimic for 6 days, and stimulated with 5uM RSG and 0.1ug/ml LPS for 24 hours. Gene expression was normalised to GAPDH. (One representative experiment out of three). *p < 0.05, **p < 0.01, ****p < 0.0001.

Several studies indicate that RSG treatment affects DC maturation and downstream activation. In line with this, we show that RSG is able to downregulate LPS induced-expression of CD83 but not CD86 in un-transfected DCs. However, in miR-511-3p^hi^ DCs, stimulation with the PPARγ antagonist GW9662 was able to significantly downregulate RSG/LPS-induced CD86 expression, most likely by reversing the effect of RSG treatment, compared to that in miR-511-3p^low^ cells (Figure 3). Taken together, our data supports the notion that inhibition of miR-511-3p could promote PPARγ activity as well as LPS-induced DC maturation/activation.

**Figure 3:**
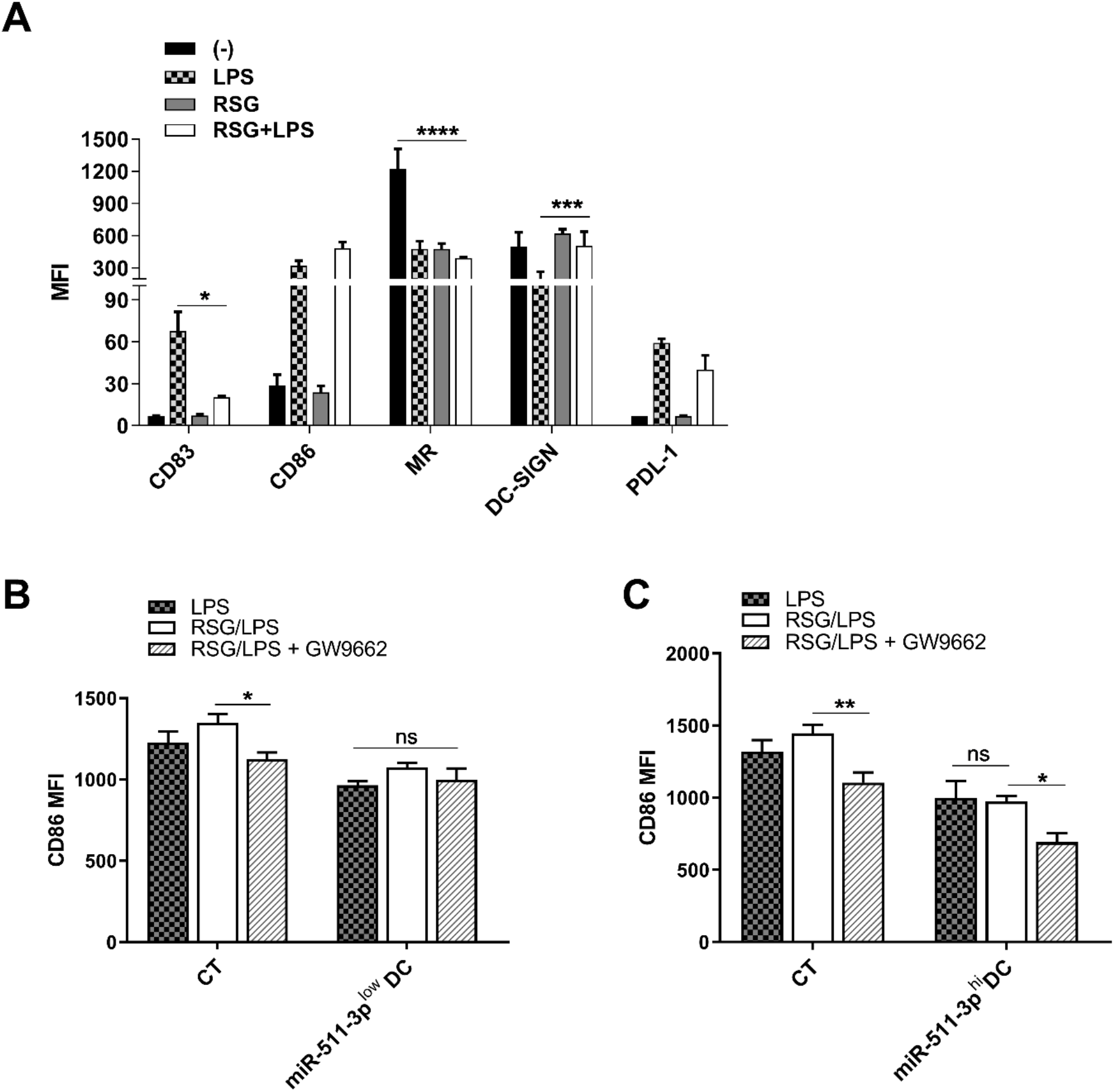
PPARγ-induced DC modulation is affected by miR-511-3p expression. A) Flow cytometry analysis of human DC markers following treatment with RSG and LPS (n=4). B and C) MFI ratios for CD86 in human miR-511-3plow and miR-5113phi DCs respectively. Monocyte derived DCs were transfected with 50nM miRNA inhibitor and mimic and stimulated with 0.1μg/ml LPS and 5μM RSG in the presence or absence of 5μM GW9662 for 24 hours (n=3). *p < 0.05, **p < 0.01, ***p < 0.001, ****p < 0.0001.

### PPARγ activation modulates cytokine production in miR-511-3p transfected DCs

It has been previously described that PPARγ activation is able to modulate LPS-induced cytokine production in cells such as macrophages and DCs from humans and mice (27–29). Having demonstrated the influence of miR-511-3p expression on PPARγ activity in human DCs, the levels of IL-6 and IL-10, pro and anti-inflammatory cytokines respectively, produced by miR-511-3p^low^ and miR-511-3p^hi^ DCs was examined to determine whether this was influenced by PPARγ activation. miR-511-3p^low^ and miR-511-3p^hi^ DCs were treated with RSG, LPS or both for 24 hours in the presence or absence of PPARγ antagonist GW9662. As shown in Figure 4, IL-6 production in miR-511-3p^low^ DCs was significantly downregulated in RSG/LPS conditions compared to LPS alone, however in the presence of the PPARγ antagonist, IL-6 was significantly increased, reverting the suppressive influence of RSG in these conditions (Figure 4A). In untransfected DCs, we found that RSG-mediated suppression of IL-6 and TNF-α production (induced by LPS treatment), was also increased after treatment with GW9662, however this was not significant (Figure S2). Conversely, miR-511-3p^hi^ DCs showed a significant increase in IL-6 after LPS treatment alone, whereas no changes were seen after GW9662 treatment. The down regulation in PPARγ expression and activity resulting from miR-511-3p overexpression could account for an increase in IL-6 production, suggesting that PPARγ could be potent suppressors of LPS-induced inflammatory responses. Interestingly, the production of IL-10 was increased in all cases following knockdown of miR-511-3p (i.e. miR-511-3p^low^ DCs) compared to CT, with exception of RSG/LPS condition, which showed no statistically significant difference with CT samples (Figure 4B). In contrast, the presence of miR-511-3p mimic treatment (i.e. miR-511-3p^hi^ DCs) significantly downregulated IL-10 production in DCs following treatment with LPS and/or RSG, which further highlights the ability of miR-511-3p overexpression to suppress the anti-inflammatory effect of PPARγ in human DCs.

**Figure 4:**
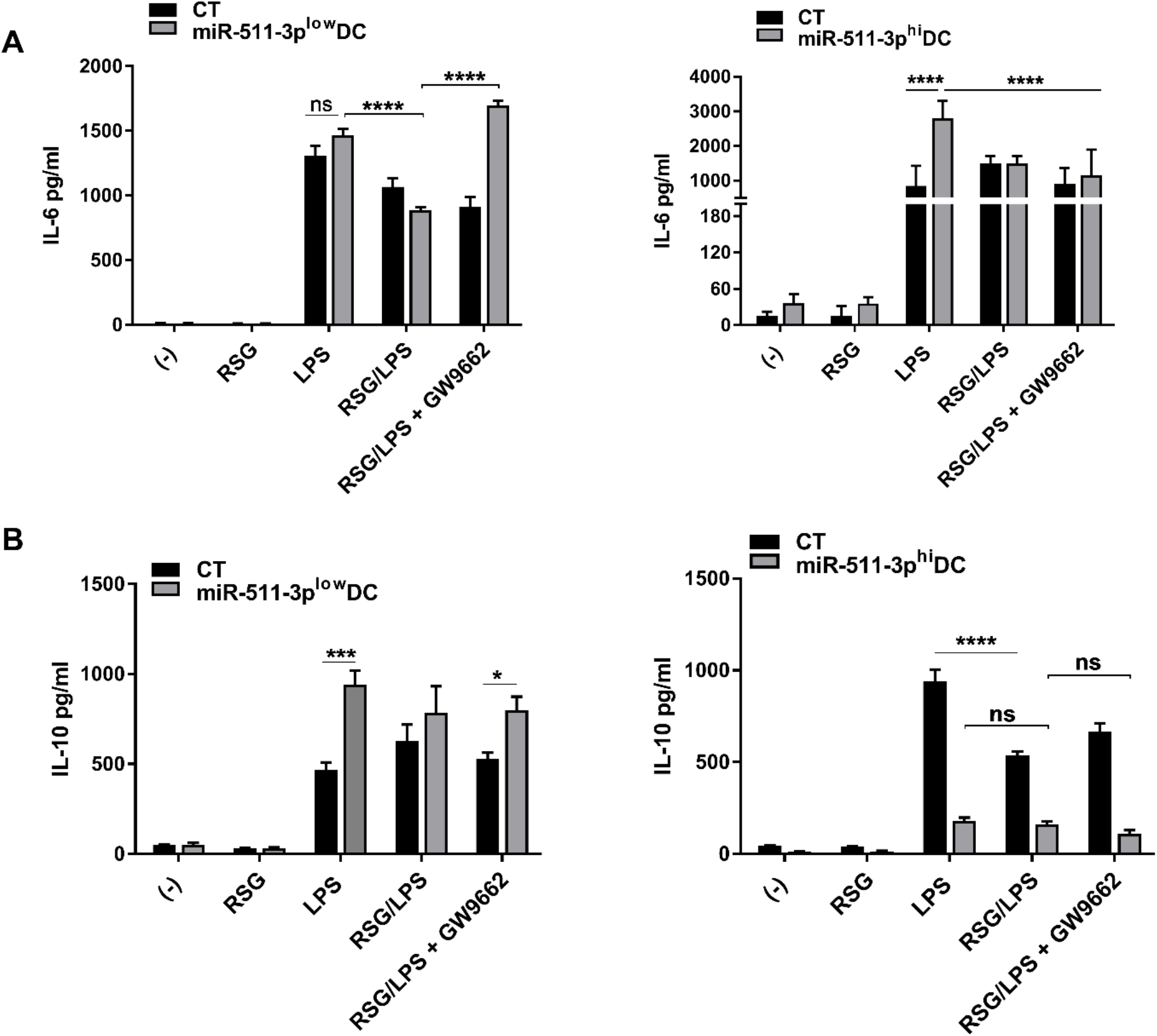
Cytokine secretion is modulated by PPARγ activity in miR-511-3p transfected DCs. A) IL-6 and B) IL-10 cytokine production by miR-511-3p^low^ and miR-511-3p^hi^ DCs respectively stimulated with 5μM RSG and 0.1μg/ml LPS for 24 hours in the presence or absence of GW9662. Culture supernatants were collected and cytokine levels were quantified by ELISA (n=3). *p < 0.05, ***p < 0.001, ****p < 0.0001.

### PPARγ modulation of LPS-induced IDO activity is influenced by miR-511-3p expression in human DCs

Under steady state conditions, induction of the tryptophan (TRP) metabolizing pathway by IDO acts as an immune regulatory mechanism in response to potent pro-inflammatory stimuli such as IFN-γ and TNF-α (30,31). Under certain conditions, IDO induction can also be mediated through an IFN-γ-independent pathway, such as presence of LPS during bacterial infection (10,32). Recently, PPARγ was also shown to promote IDO activity and generation of local tolerogenic DCs within solid tumours (14), nonetheless the link between PPARγ and IDO in DCs within the context of LPS-induced inflammation is unknown. In order to investigate this, we first treated untransfected DCs with LPS, RSG and GW9662 alone to determine the effect of these ligands on IDO activity (Figure S3). Subsequently, we determined IDO activity following treatment with LPS in the presence of these ligands. IDO activity was measured by colorimetric determination of kynurenine (KYN) produced in culture supernatant. As summarised in Figure 5, RSG alone had no influence on IDO activity, but significantly downregulated LPS-induced IDO activity in DCs. Additionally, treatment with the PPARγ antagonist (RSG/LPS + GW9662) caused a decrease in IDO activity compared with RSG/LPS condition alone, however this was not statistically significant (Figure 5A). Given the effect of miR-511-3p on PPARγ expression and activity, we then examined whether changes in miR-511-3p expression in DCs could affect IDO activity following treatment with LPS and RSG. Again, RSG stimulation alone had no effect on IDO activity (data not shown), but was able to significantly downregulate LPS-induced IDO activity in CT and miR-511-3p^low^ DCs (Figure 5B). Interestingly, IDO activity was markedly reduced in the miR-511-3p^hi^ cells after treatment with RSG and LPS similar to observations with IL-10. Furthermore, we found an inverse correlation between miR-511-3p regulation and IDO activity after treatment with GW9662, indicating that PPARγ may play a key role in IDO downregulation induced by RSG in human DCs (Figure 5C). These data suggest that PPARγ activation is able to downregulate IDO activity in human DCs partly due to changes in miR-511-3p expression. As indicated in the LIVE/DEAD stain, DC viability was comparable in all conditions tested (Figure S4).

**Figure 5:**
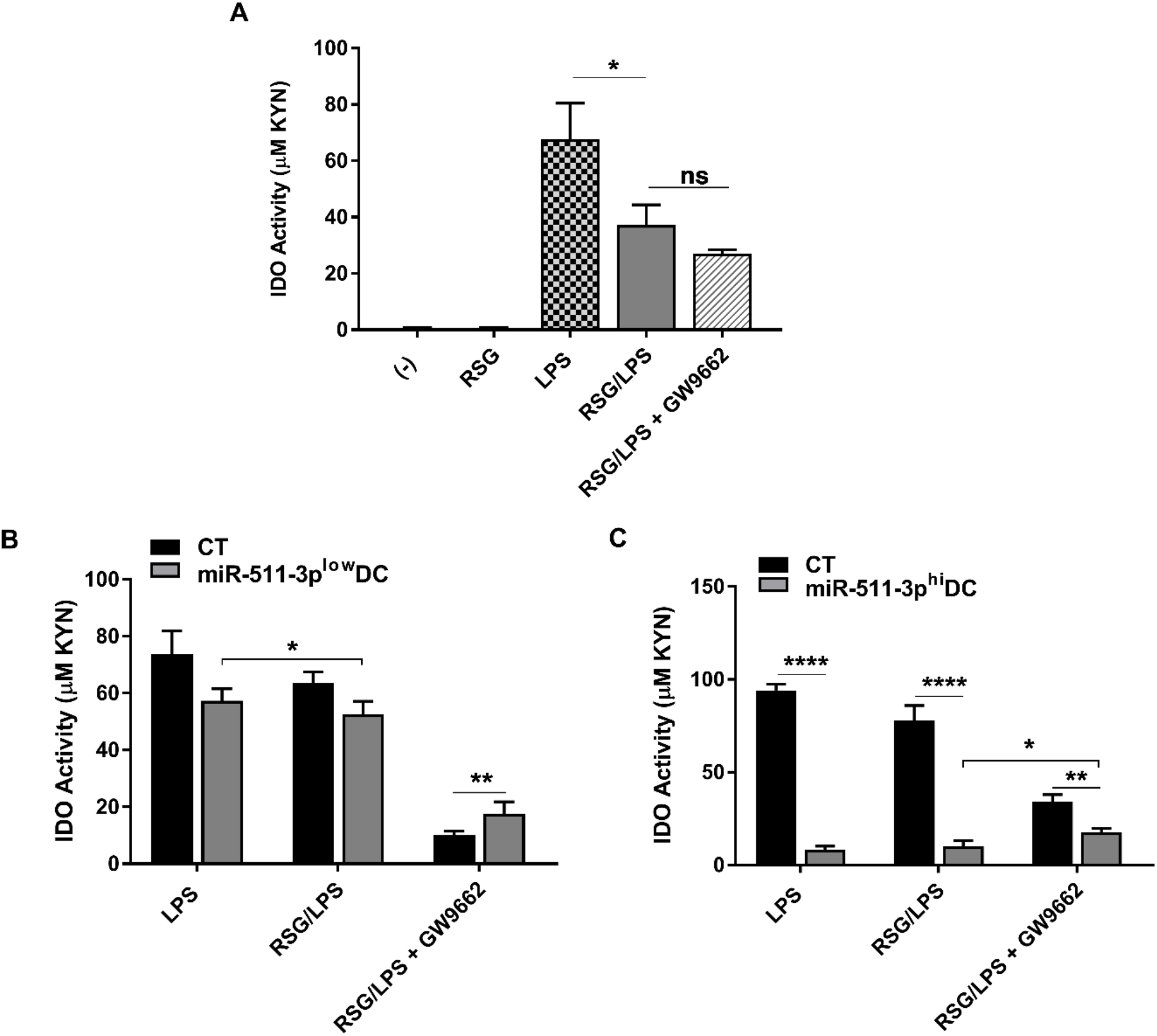
IDO regulation by human monocyte-derived dendritic cells. A) IDO activity in human DCs treated with 5μM RSG and 0.1μg/ml LPS for 24 hours in the presence or absence of 5μM GW9662 (n=3). B and C) IDO activity in miR-511-3p^low^ and miR-511-3p^hi^ DCs respectively following treatment with 5μM RSG and 0.1μg/ml LPS for 24 hours in the presence or absence of 5μM GW9662. Culture supernatant was collected and examined for IDO activity (n=3). *p < 0.05, **p < 0.01, ****p < 0.0001.

## Discussion

The PPAR receptors, which belong to the nuclear hormone receptor superfamily, were originally identified as key players in controlling oxidation of lipids and fatty acids. Among the three members, PPARγ is essential for controlling adipocyte differentiation and glucose metabolism (33) and presently, several reports have demonstrated the importance of PPARγ agonists as treatment against inflammatory diseases (34–36). MicroRNAs are crucial players in mammalian gene regulation and their role in fine-tuning gene expression, particularly through dysregulated expression has been reported in several infectious and inflammatory diseases. In this study, the role of miR-511-3p in modulating PPARγ activity in human DCs was investigated within the context of LPS-induced inflammatory responses.

Tserel et al, using TargetScan and a combination of different algorithms highlighted *PPARG* as a putative target of miR-511 activity (16). In line with this, data presented in Figure 1 show that overexpression of miR-511-3p significantly downregulated PPARγ expression as quantified by qRT-PCR. Recently, miRNAs have been shown to mediate pathways decreasing PPARγ expression during the onset of inflammation. For instance miR-27b was shown to decrease PPARγ mRNA level, as mutation or deletion of the miR-27b start site completely abolished reduction of luciferase activity in THP1 macrophages (37). Additionally, they showed that miR-27b inhibitors prevented LPS-dependent PPARγ mRNA reduction in a concentration-dependent manner.

In line with previous reports, our data show that treatment of DCs with the PPARγ agonist rosiglitazone (RSG), induced PPARγ activity, whereas LPS treatment alone completely abrogates PPARγ activity (Figure 2). Two isoforms of PPARγ have been identified (PPARγ1 and 2) and studies show that both isoforms are downregulated in monocyte and macrophage cell lines following treatment with LPS (37). However, prolonged exposure to LPS in these studies enabled recovery of PPARγ mRNA levels to almost basal levels after 24 hours (37).

Notably, Samokhvalov et el demonstrated that although LPS downregulates PPARγ activity, epoxyeicosatrienoic acids (EETs), which are biologically active metabolites of arachidonic acids, act as PPARγ agonists to suppress LPS-induced pro-inflammatory responses (38). This could explain the increase in PPARγ activity following RSG+LPS treatment in miR-511-3p^low^ DCs seen in the present study (Figure 2A). Similarly, in a rat model of sepsis, infusion of low-dose LPS was shown to significantly decrease hepatic PPARγ protein levels, but administration of the LPS binding agent (polymyxin B) reduced plasma endotoxin level and prevented PPARγ downregulation in the septic animals (39), thus highlighting the complex mechanisms underlying the regulation of PPARγ during LPS-induced inflammation.

Several reports have indicated that induction of the IDO pathway of tryptophan metabolism contributes towards immune modulatory events (40). Within the context of inflammation, IDO induction leads to the suppression of T cell effector responses (41). A role for LPS in IDO induction has also been demonstrated by our group and others (19,42), which explains why IDO activity and expression is increased following LPS treatment. In this study, we show that LPS induced IDO activity is significantly reduced in the presence of RSG compared to LPS alone. Interestingly, IDO activity following treatment with RSG and/or LPS in miR-511-3p^hi^ DCs was also significantly reduced when compared to CT cells only. It is likely that transfection with miRNA mimics resulted in repression of other cell responses or mRNA targets as previously reported (43). This could account for the decrease in IDO activity in miR-511-3p^hi^ cells. Recently, it has been demonstrated that fatty acid oxidization by PPARγ in melanomas positively regulates IDO in DCs, by promoting a switch towards a tolerogenic state to enable immune evasion (14). In contrast, our data indicate that PPARγ activation may negatively regulate IDO activity in DCs within the context of LPS-induced inflammation. A schematic representation of the relationship between miR-511-3p and PPARγ in modulating DC function is shown in Figure 6.

**Figure 6:**
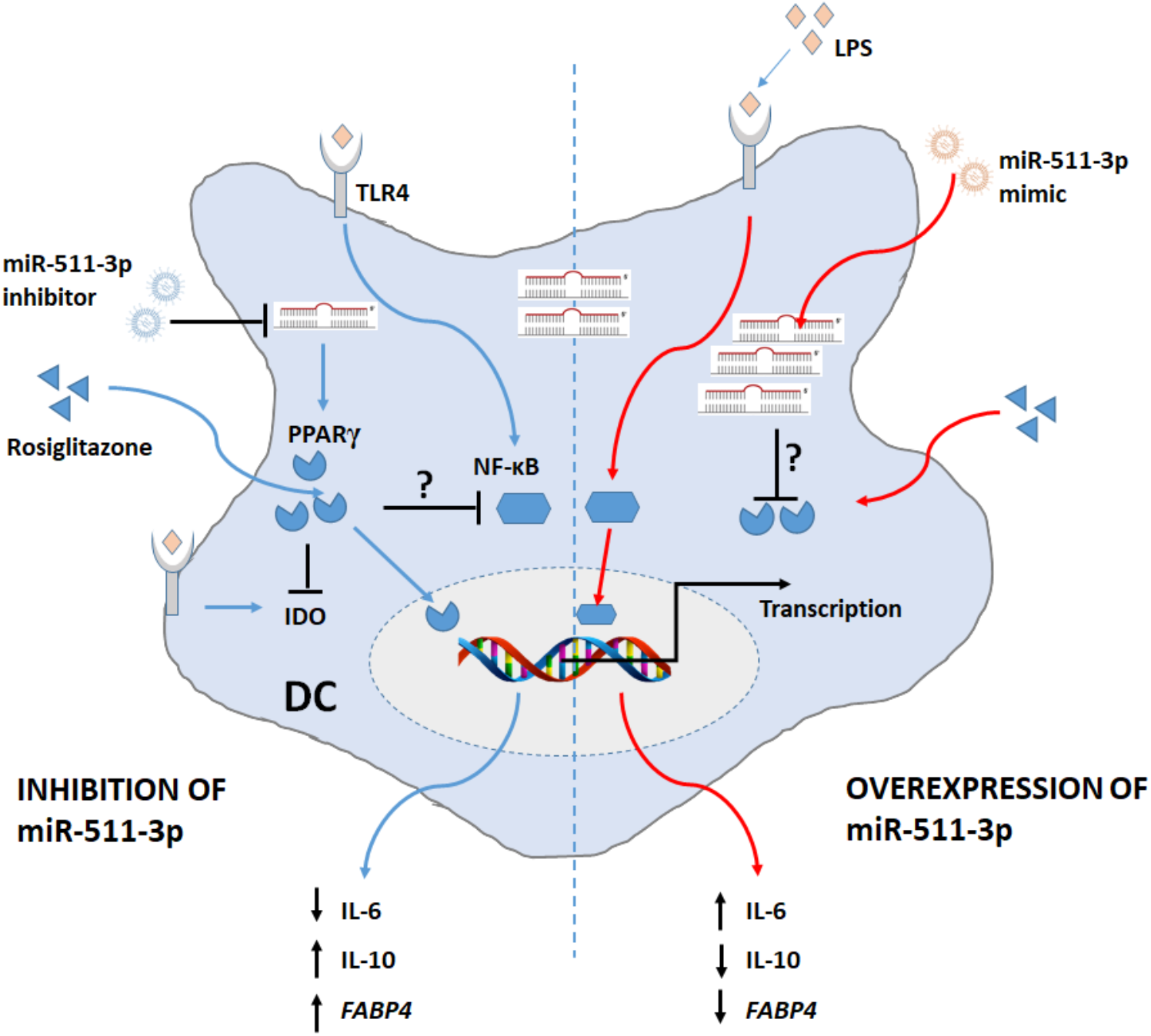
Schematic representation of the relationship between miR-511-3p and PPARγ in modulating human DCs function. Downregulation of miR-511-3p promotes of PPARγ expression and activity (*FABP4*) and subsequently suppresses LPS-induced inflammation most likely through inhibiting NF-κB. In contrast, overexpressing miR-511-3p reverses the effects of PPARγ and promotes transcription of pro-inflammatory genes.

The anti-inflammatory effect of PPARγ ligands in regulating immune responses is well documented. In particular, high dose 15d-PGJ2 or TZD (PPARγ agonists) treatment of monocytes and macrophages was shown to inhibit secretion of pro-inflammatory cytokines (IL-6, TNF-α and IL-1β) (26), and suppress IFN-γ-dependent inducible genes (25,44). Additionally, it has been shown that IL-10 is induced by RSG in experimental models of colitis and Parkinson’s disease (45,46). In the present study, we highlight the dynamics of IL-6 and IL-10 production by miR-511-3p modified DCs, which further demonstrates the role of PPARγ supporting an anti-inflammatory profile in miR-511-3p^low^ DCs. Interestingly, a number of potential links between PPARγ and c-type lectins, particularly the mannose receptor (MR) have also been suggested. These include data showing that interaction between MR and Man-LAM from *Mycobacterium tuberculosis* can induce PPARγ activation (47). Considering that changes in miR-511-3p expression is associated with changes in MR expression as well as IDO activity as demonstrated previously (24), it is likely that miR-511-3p may play a role in regulating MR expression and DC function through PPARγ. These studies therefore suggest that PPARγ and signalling molecules/pathways controlling its expression or activity may serve as a target for anti-inflammatory therapy. Collectively, these data highlight the potential immune-regulatory role of miR-511-3p on PPARγ activity in human DCs. The roles of miRNAs and PPARγ in immune regulation are still being investigated and this transcription factor is emerging as a key player in various stages of the resolution of inflammation.

## Acknowledgements

Authors would like to acknowledge support from the flow cytometry facility (School of Life Sciences, University of Nottingham). D. Awuah is a recipient of the Vice Chancellor’s Ph.D. Scholarship at the University of Nottingham, United Kingdom.

## Author Contributions

D.A designed and carried out experimental work and wrote the manuscript. A.R, M.A and C.M carried out experimental work. A.M.G. designed and supervised the work, and revised the manuscript.

## Abbreviations

CT: Scrambled control
DC: Dendritic cell
FABP4: Fatty acid binding protein 4
IDO: Indoleamine 2,3-dioxygenase
LPS: Lipopolysaccharide
MFI: Median fluorescence intensity
PPARγ: Peroxisome proliferator-activated receptor gamma
RSG: Rosiglitazone
TLR: Toll-like receptor

