## Supplemental information for "MicroRNA-511-3p mediated modulation of the peroxisome proliferator-activated receptor gamma (PPARγ) controls LPS-induced inflammatory responses in human monocyte derived DCs"

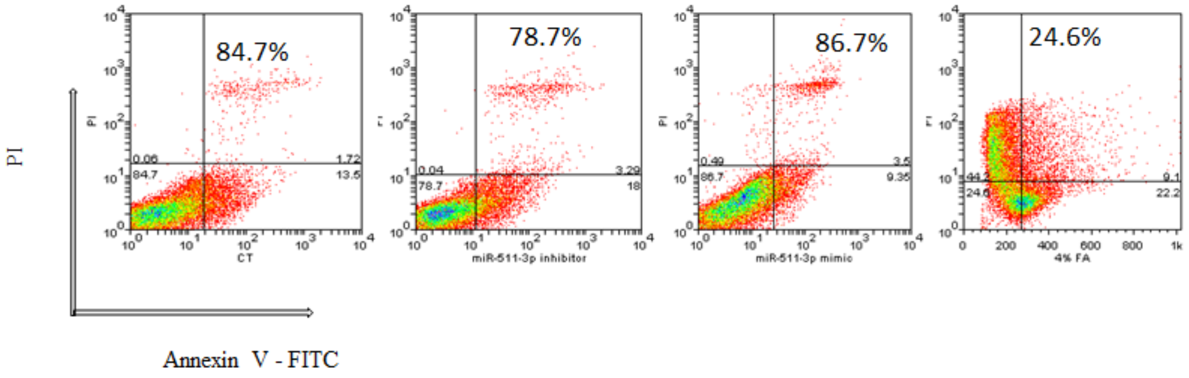


**Supplementary Figure 1: Examining apoptotic effect of miR-511-3p transfection on Mo-DCs.** Representative dot plots are shown for monocyte-derived DCs transfected with miR-511-3p inhibitors and mimics and harvested on day 6 for viability analysis with Annexin V apoptosis detection kit. Panels represent unstained plots for scrambled control (CT), inhibitor and mimic treatment, and formaldehyde treated positive control. Cells were analysed by flow cytometry within 30 minutes of staining. Representative data shown is from one independent experiment with a single donor out of a total of two donors.

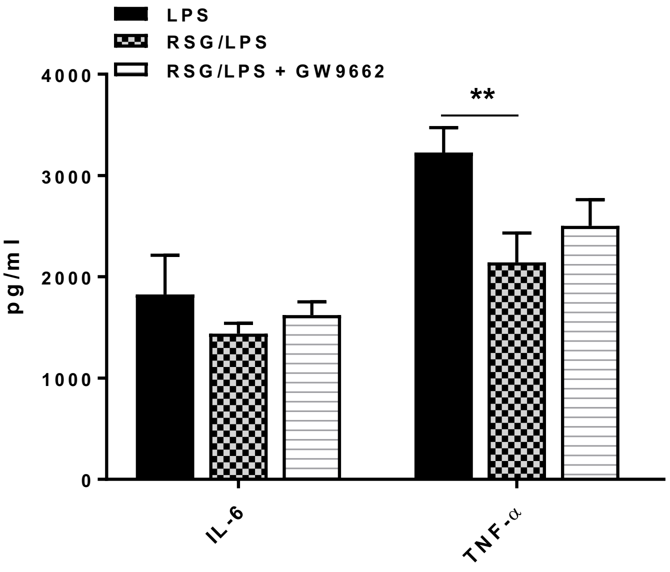


**Supplementary Figure 2: PPARγ activation by rosiglitazone (RSG) downregulates pro-inflammatory cytokine production in human DC**. LPS induced TNF-α production is significantly suppressed by RSG which is partially reverted in the presence of GW9662, a PPARγ antagonist (n=4). **p < 0.01.

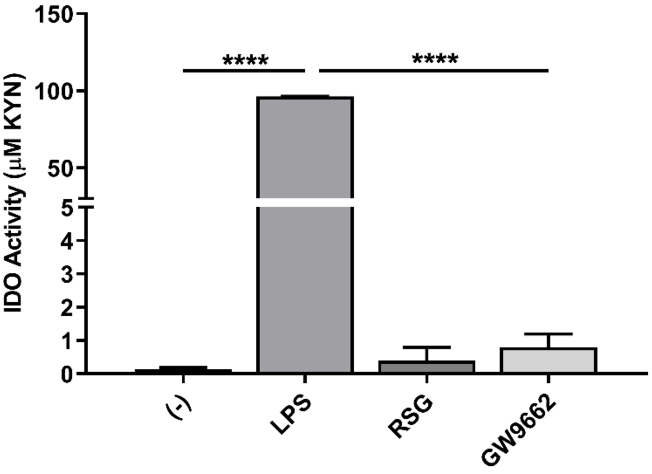


**Supplementary Figure 3: IDO activity in un-transfected DCs**. Monocyte derived DCs were treated with 100ng/ml of LPS, 10uM of RSG and GW9662 for 24 hours. Culture supernatants were harvested and IDO activity was measured afterwards (n=3). ****p < 0.0001.

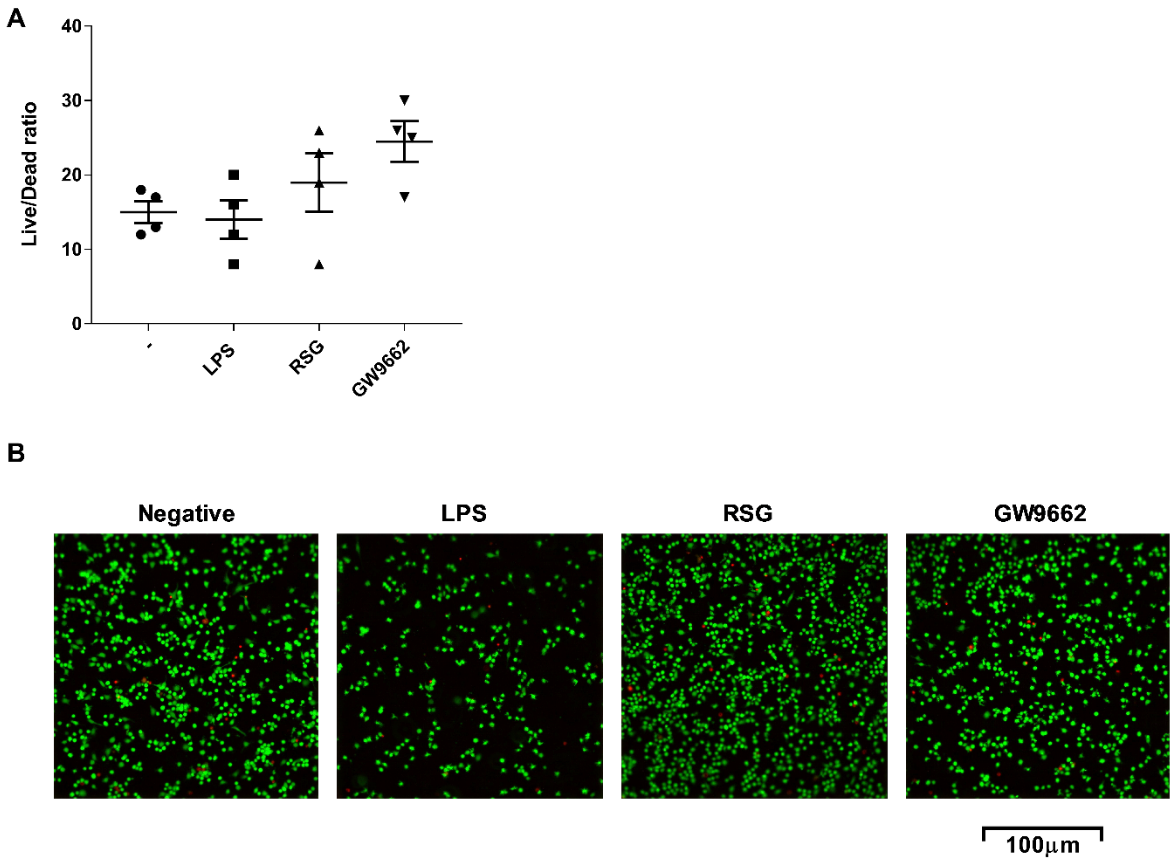


**Supplementary Figure 4**: **Dendritic cell viability** A and B) Live/Dead ratios for cell viability following treatment with LPS, RSG and GW9662. Live and dead cells are represented by green and red fluorescence respectively. Images are representative of 2 independent experiments.
